# Amphiregulin mediates non-cell-autonomous effect of senescence on reprogramming

**DOI:** 10.1101/2021.09.01.458621

**Authors:** Mathieu von Joest, Cheng Chen, Thibaut Douché, Aurelie Chiche, Mariette Matondo, Han Li

**Affiliations:** Cellular Plasticity & Disease Modelling, Dept. of Developmental & Stem Cell Biology, CNRS UMR 3738; Proteomics Platform, Mass Spectrometry for Biology Unit (MSBio), USR CNRS 2000, Institut Pasteur, 25 rue du Dr Roux, Paris 75015, France; Laboratory of chromatin and gene regulation during development, Imagine Institute, INSERM UMR1163, 75015, Paris, France

**Keywords:** cellular senescence, SASP, in vitro & in vivo reprogramming, cellular plasticity, amphiregulin

## Abstract

Cellular senescence is an irreversible growth arrest with a highly dynamic secretome, termed the senescence-associated secretory phenotype (SASP). Senescence has been implicated in somatic reprogramming to pluripotency. The cell-intrinsic proliferation arrest is a barrier for reprogramming, whereas the SASP facilitates the cell fate conversion in nonsenescent cells. However, the mechanisms by which reprogramming-induced senescence regulates cell plasticity are not well understood. Here, we have further investigated how the heterogeneity of paracrine senescence impacts reprogramming. We show that senescence promotes *in vitro* reprogramming in a stress-dependent manner. We identified a catalog of SASP factors and pathways potentially involved in the cell fate conversion using an unbiased proteomic analysis. Amphiregulin (AREG), a growth factor frequently secreted by the senescent cells, promotes *in vitro* reprogramming by accelerating proliferation and MET via the EGFR signaling pathway. Of note, AREG treatment diminished the negative effect of donor age on reprogramming. Finally, AREG enhances *in vivo* reprogramming in the skeletal muscle. Hence, senescence could facilitate cellular plasticity via various SASP factors to promote reprogramming and tissue repair.

## BACKGROUND

Cellular senescence is a stress response characterized by a permanent cell-cycle arrest, acquisition of a senescence-associated secretory phenotype (SASP), and resistance to apoptosis (Hernandez-Segura et al., 2018). Senescence is involved in variety biological and pathological processes including development, tissue repair, tumorigenesis, and organism aging (Munoz-Espin and Serrano, 2014; Rhinn et al., 2019). Of note, SASP plays a crucial role in mediating senescence non-cell autonomous functions by interacting with neighboring cells and the surrounding microenvironment (Ito et al., 2017). It is becoming clear that senescence phenotypes, particularly SASP composition, are highly heterogeneous and depending on the initial stimuli, the affected cell type, and the anatomic location (Hernandez-Segura et al., 2017) (Ito et al., 2017).

Cellular reprogramming is a process of converting fully differentiated cells back to the pluripotent state, which has tremendous potentials in many areas of biomedical research (Takahashi and Yamanaka, 2006; Robinton and Daley, 2012). Interestingly, recent studies revealed that senescence has both cell-autonomous and non-cell-autonomous effects on reprogramming (Aarts et al., 2017; Banito et al., 2009; Chiche et al., 2017; Mosteiro et al., 2016; Mosteiro et al., 2018). Ectopic expression of reprogramming factors: Oct4, Klf4, Sox2, and c-Myc (OSKM), induced senescence as a cell intrinsic barrier for reprogramming (Banito et al., 2009), which in turn promote cellular plasticity and reprogramming in the non-senescent cells via SASP, in particularly by interleukin-6 (IL-6) (Chiche et al., 2017; Mosteiro et al., 2016; Mosteiro et al., 2018). However, the mechanisms by which paracrine senescence, with specific SASP factor(s), regulate cellular plasticity in the context of reprogramming remain not fully understood.

The epidermal growth factor receptor (EGFR) signaling pathway is one of the most important pathways that regulate cellular growth, proliferation, differentiation, and survival (Herbst, 2004). Although it is important for embryonic stem cells functions (Schuldiner et al., 2000; Yu et al., 2019) and the early embryonic development (Kim et al., 1999), the role of the EGFR signaling pathway in somatic reprogramming remains poorly understood. A variety of senescent cells produce abundant amphiregulin (AREG) (Coppe et al., 2010), a low-affinity EGFR ligand (Jones et al., 1999; Shoyab et al., 1988). Of note, accumulating evidence indicates AREG is a vital mediator of the immune response in infection and tissue repair (Zaiss et al., 2015). It has been suggested that AREG regulates senescence (Pommer et al., 2021) and contributes to an immunosuppressive tumor microenvironment (Xu et al., 2019). Given the pleiotropic functions of AREG, its roles in mediating senescence functions remain largely unexplored.

In this study, we investigated how paracrine senescence impacts reprogramming focusing on the stress dependent SASP heterogeneity and identified a collection of SASP factors potentially important for cellular plasticity using unbiased proteomic analysis. Next, we demonstrated that AREG enhances reprogramming both *in vitro* and *in vivo* via EGFR and alleviates the age-associated reduction in reprogramming.

## RESULTS

### Paracrine senescence promotes *in vitro* reprogramming in a stress-dependent manner

Given senescence can impact reprogramming in both cell-autonomous and non-cell-autonomous manner (Chiche et al., 2020), and SASP composition is highly context-dependent, determined by both the initial stress and the tissue/cell of origin(Hernandez-Segura et al., 2017; Ito et al., 2017). We reasoned stress dependent SASP might impact reprogramming efficiency differently. To address this question, we treated non-reprogrammable wild-type (WT) primary mouse embryonic fibroblasts (MEFs) with three different stimuli (replicative senescence (RS), γ-irradiation (IRIS), and oncogene-induced senescence (OIS)) to induce senescence and generate distinct SASP (Figure S1A). Meanwhile, we used the MEFs generated from a previously established reprogrammable mouse model (i4F) (Abad et al., 2013). The reprogramming was induced by adding doxycycline (DOX) in i4F-MEFs either co-cultured with the senescent WT MEFs (co-culture) or cultured directly with the senescence conditioned medium (CM) according to the experimental scheme illustrated (Figure 1A). After 12 days DOX treatment, induced pluripotent stem cells (iPSCs) colonies were stained using alkaline phosphatase (AP) to determine the reprogramming efficiency. For the co-culture system, i4F MEFs were seeded on top of 2×10^5^ WT MEFs (either control or OIS treated) with various ratios of 1:3, 1:5, and 1:10 (i4F: WT), to determine the optimal experimental condition. We found the reprogramming efficiency increased the most in the ratio of 1:10 (i4F: WT) (Figure S1B). Next, we compared the relative reprogramming efficiency of i4F MEFs and found that in both systems all stimuli can increase reprogramming efficiency ranging from RS the least to OIS the most, except the positive effect of RS is diminished in the CM system (Figure 1B & Figure 1C).

**Figure 1.**
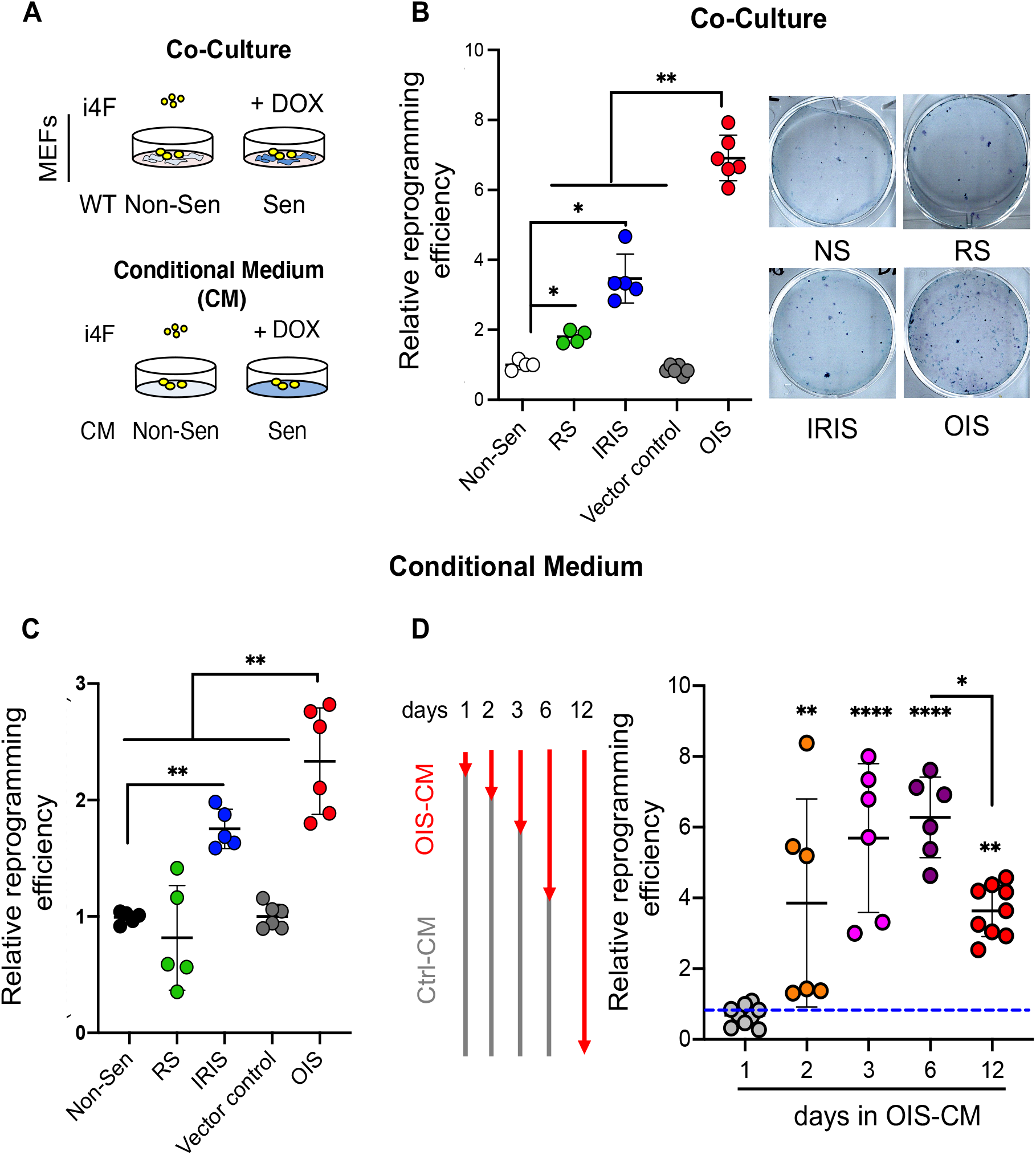
Senescence promotes *in vitro* reprogramming in a stress-dependent manner. **A.** Scheme of the experiments. **B & C.** Relative *in vitro* reprogramming efficiency of i4F MEFs exposed to secretome of non-senescent compared to senescent cells generated by different stress (corresponding to the respective controls: RS and IRIS to non-sen and OIS to vector alone). RS: replicative senescence, IRIS: γ-irradiation, OIS: oncogene-induced senescence. **B.** Co-culture system; Representative image of plate stained with alkaline phosphatase (AP) to reveal colonies arising from reprogramming (right panel). **C.** Conditioned medium system (CM). **D.** Reprogramming fold change of i4F MEFs cultured in different duration of OIS-CM before switching to Ctrl-CM. Scheme of the experiment (left panel). All the reprogramming fold changes was normalized to i4F in Ctrl-CM for 12 days, indicated by the blue dash line. Data correspond to the average ± s.d. using indicated n number of independent experiments. Statistical significance was assessed by ordinary one-way ANOVA test: *p < 0.05; **p < 0.01; **** p < 0.0001. See also **Figure S1**.

Reprogramming is a stepwise process with distinct phases: initiation, maturation, and stabilization (Samavarchi-Tehrani et al., 2010). To determine at which stage the SASP exposure is most important for enhancing reprogramming, we cultured i4F with either OIS-CM or Ctrl-CM in different order (Figure 1D & Figure S1C). Culturing i4F-MEFs with OIS-CM for the first three days was sufficient to induce the highest fold increase in reprogramming (Figure 1D). Conversely, initiating reprogramming with Ctrl-CM for three days and replaced by OIS-CM completely abolished the beneficial effect (Figure S1C). Active cell proliferation is rate-limiting for successful reprogramming (Ruiz et al., 2011), and senescent cells secrete various growth factors (Coppe et al., 2010). Therefore, we compared the growth properties of i4F MEFs cultured with either OIS-CM or Ctrl-CM over 12 days in the absence of DOX. The bromodeoxyuridine (BrdU) incorporation rates after 72 hours were similar between Ctrl and OIS (Figure S1E). However, the accumulative growth curve indicated that i4F MEFs grew faster in OIS-CM than in Ctrl-CM (Figure S1D). Hence, certain SASP factors promote *in vitro* reprogramming transiently at the initiation phase of reprogramming, which partially correlates with an enhanced proliferation rate.

### Conditioned medium of OIS-Il6^-/-^ promoted reprogramming efficiently

We sought to identify SASP factors responsible for the stress-dependent effect on reprogramming. Given that IL-6 signaling is required for *in vitro* reprogramming (Brady et al., 2013; Mai et al., 2018), and IL-6 is a crucial mediator of the senescence paracrine signals in promoting *in vivo* reprogramming (Chiche et al., 2017; Mosteiro et al., 2016; Mosteiro et al., 2018), we determined firstly the effect of CM of senescent MEFs derived from *Il6^-/-^* mice on reprogramming. Of note, IL-6 plays a critical role in mediating OIS, and reduction of IL-6 via RNA interference efficiently bypassed OIS in human diploid fibroblasts (Kuilman et al., 2008). Interestingly, we found *Il6^-/-^* MEF entered senescence efficiently upon OIS, judging by both senescence-associated beta-galactosidase (SAβGal) staining (data not shown) and qPCR (Figure S2A). Surprisingly, CM generated from senescent *Il6^-/-^* MEFs by IRIS or OIS enhanced reprogramming efficiency to a similar level as WT (Figure 2A). To ensure the observed enhancement is IL-6 independent, we compared the IL-6 level in the OIS-CMs of both WT and *Il6^-/-^* by ELISA. In WT, IL-6 is significantly increased in the OIS-CM compared to Ctrl-CM. While, IL-6 is non-detectable in both Ctrl-CM and OIS-CM of *Il6^-/-^* (Figure S2B). Of note, IL-6 neutralizing antibody could potently repress the increased reprogramming efficiency only in the CM-WT but failed to do so in the CM-*Il6^-/-^* (Figure 2B and Figure S2C). Thus, there are SASP factors other than IL-6 that can robustly promote *in vitro* reprogramming.

**Figure 2.**
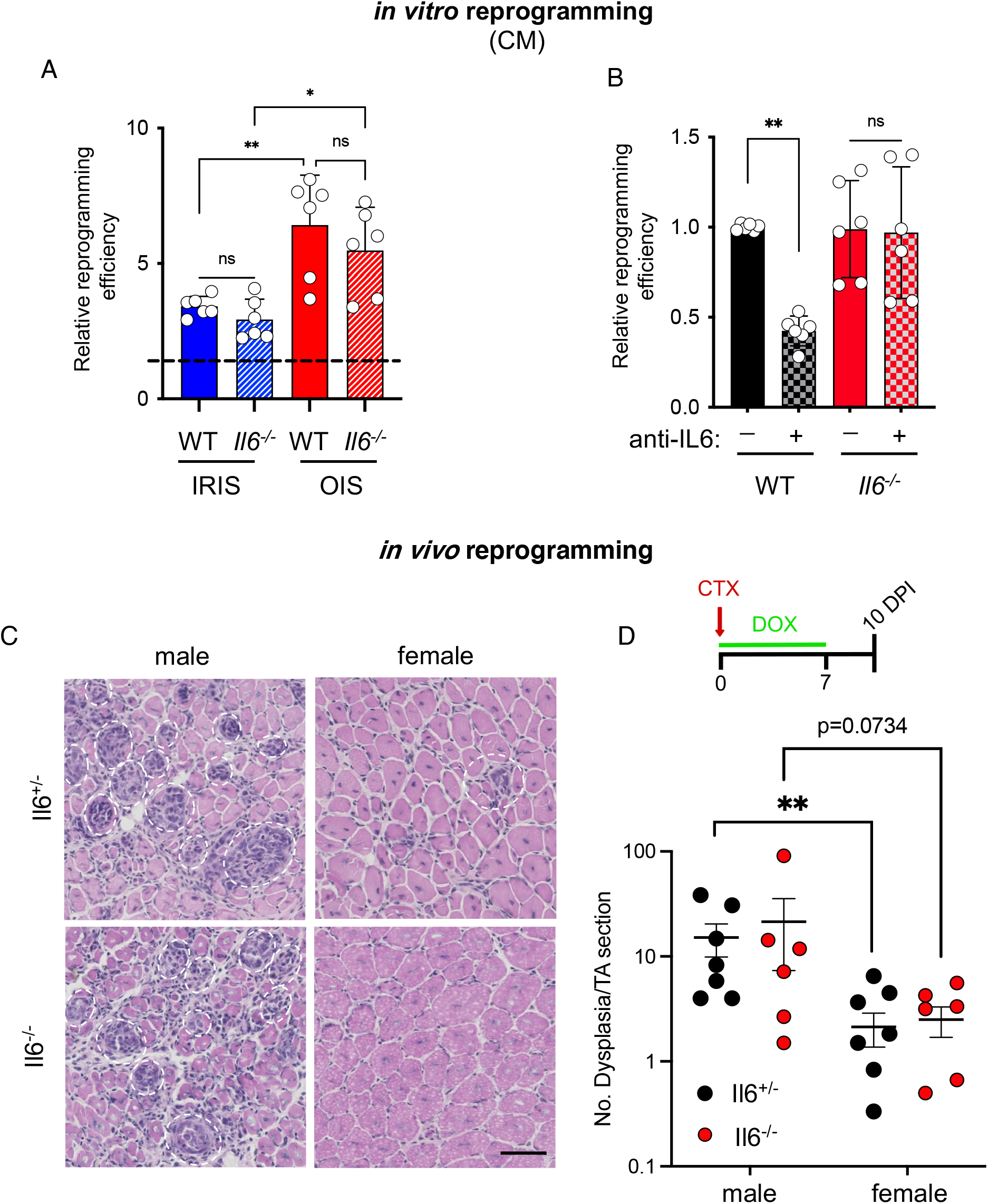
*in vivo* reprogramming in the skeletal muscle occurs normally in *Il6^-/-^* mice. **A.** Relative reprogramming efficiency of the i4F MEFs cultured in the CM derived from either WT or *Il6^-/-^* senescent cells. Senescence was induced using two different stresses, IRIS and OIS. Fold change was normalized with CM derived of respective non-senescent counterparts. **B.** Reprogramming efficiency of i4F MEFs cultured in CM of WT or *Il6^-/-^* senescent cells compared to treatment of IL-6 blocking antibody. Senescence was induced by over-expression of oncogene (OIS). Data correspond to the average ± s.d. using indicated n number of independent experiments. **C.** Transverse TA cryosections of indicated genotypes and sex after histological staining with Haematoxylin and Eosin (H&E). Dashed circle highlights dysplasia region. Scale bar: 50 μm. **D.** Experimental scheme (Upper panel) and quantification of **C.** The value represents one TA per mouse. Statistical significance was assessed by ordinary one-way ANOVA test: *p < 0.05; **p < 0.01; ***p < 0.001. See also **Figure S2**.

### *In vivo* reprogramming in the skeletal muscle is not affected by IL6 deficiency

We reported previously that injury-induced senescence enabled reprogramming in the skeletal muscle (Chiche et al., 2017). We wondered whether tissue injury could induce reprogramming in the skeletal muscle of *Il6^-/-^* mice. To test this, we generated *Il6^-/-^*;i4F mouse model and injured the Tibialis anterior (TA) muscle of both *Il6^+/-^*;i4F and *Il6^-/-^*;i4F mice with the snake venom cardiotoxin (CTX). In the same time, we added doxycycline (dox) in the drinking water for 7 days to induce *in vivo* reprogramming and TAs were collected at 10 days post-injury (DPI) as previously described (Chiche et al., 2017). The expression of the OSKM triggered extensive dysplasia and reprogramming (Nanog^+^ cells) in the injured TA of male i4F mice (Figure 2C and Figure S2D). Consistent with a previous report (Mosteiro et al., 2018), the reprogramming was significantly reduced in the females i4F mice comparing to male i4F mice (Figure 2C & 2D,0 Figure S2D). Interestingly, *Il6^-/-^* i4F mice exhibited a similar level of dysplasia and Nanog^+^ cells as the sex-matched *Il6^+/-^*;i4F mice (Figure 2C & 2D, Figure S2D). Besides, we did not observe a difference in the senescence induction in males regardless of the genotype (Figure S2E & S2F). Thus, *in vivo* reprogramming in the skeletal muscle is not impaired in the IL-6 deficient reprogrammable mice.

### Soluble factor fraction of the CM-OIS mediates *in vitro* reprogramming

To gain a comprehensive profile of the SASP heterogenous composition, we determined firstly the effect of different fractions of CM on reprogramming, which could facilitate reprogramming due to secreted proteins, extracellular vesicles, or small-molecule metabolites. We examined whether the boiled CM could promote *in vitro* reprogramming since boiling denatures most proteins without affecting small molecule metabolites, and found that boiled CM completely abolished *in vitro* reprogramming, indicating that secreted proteins are crucial in promoting reprogramming (Figure S3A).

Exosomes are small extracellular vesicles (sEV) released from almost all cell types and are critical mediators of intercellular communication (Kalluri and LeBleu, 2020). Increasing evidence suggests that senescent cells secret exosomes with distinct features, contributing to paracrine senescence (Borghesan et al., 2019). To examine whether senescence-associated exosomes regulate *in vitro* reprogramming, we isolated sEV and supernatants (SN) fractions from Ctrl-CM and OIS-CM using serial ultracentrifugation and filtering with a 0.22-μm filter (Thery et al., 2006). Both OIS-sEV and Ctrl-sEV were suspended in Ctrl-SN to determine whether they can drive paracrine senescence. Consistent with the previous report (Borghesan et al., 2019), the OIS-sEV in Ctrl-SN induces several senescence markers in i4F MEFs, as shown by a decrease in BrdU incorporation and an increase in mRNA expression of p16 and ARF (Figure S3B & Figure S3C). Next, we wondered whether OIS-sEV could also affect reprogramming. Comparing to Ctrl-SN, the OIS-SN significantly enhanced the reprogramming efficiency to a similar level as OIS-CM (Figure 3A). However, OIS-sEV in Ctrl-SN exhibited a much lower reprogramming efficiency than OIS-SN, similarly to Ctrl-sEV in Ctrl-SN (Figure 3A). Hence, the soluble fraction of OIS-CM is essential for promoting in vitro reprogramming.

**Figure 3.**
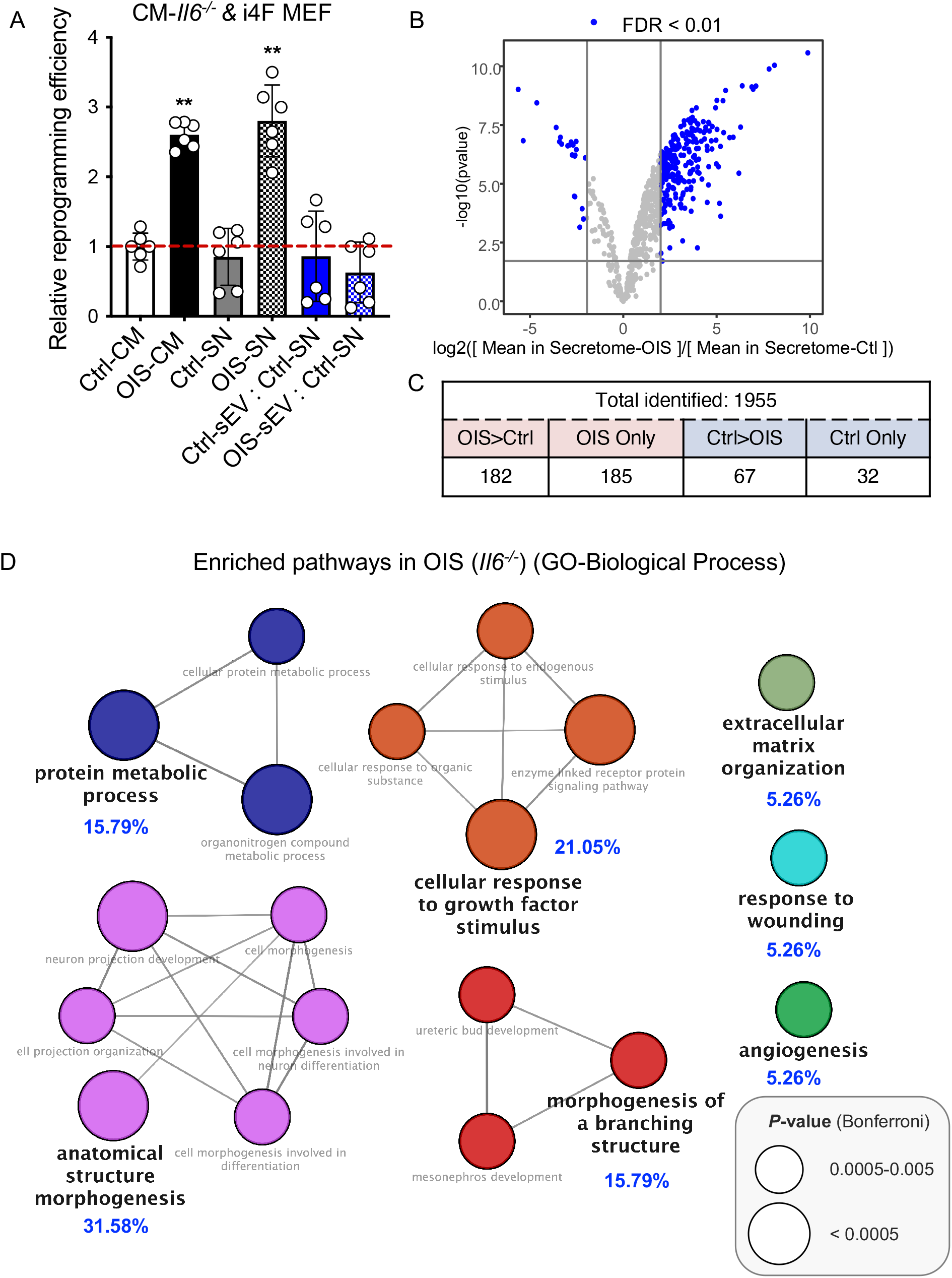
Proteomic analysis of oncogenic stress-induced secretome in *Il6^-/-^* MEFs. **A.** Fold change of reprogramming efficiency of i4F MEFs cultured in indicated different combination of various fractions of CM from non-senescent (Ctrl) or senescent (OIS) *Il6^-/-^*. SN: supernatants (soluble proteins); sEV: small extracellular vesicles (exosome). Reprogramming efficiency fold change normalized to Ctrl-CM. Data correspond to the average ± s.d. using indicated n number of independent experiments. **B.** Volcano plot shows log_2_ fold change in protein intensity against the -log10 pvalue in SEN compared to Ctrl; blue dots are proteins with significant differential expression (FDR< 0.01). **C.** Summary of proteins either with significantly altered secretion (q-value < 0.01 and >2-fold change) by senescent (SEN) compared with quiescent cells (Ctrl), or exclusively detected in all repeats from one condition, following oncogenic stress (OIS) in senescent *Il6^-/-^* primary MEFs. **D.** ClueGO (Bindea et al., 2009) pathway enrichment and network analyses of OIS-secretome. Pathways of the same color have ≥50% similarity. Connecting lines represent Kappa connectivity scores > 40%. Statistical significance was assessed by ordinary one-way ANOVA test: **p < 0.01. See also **Figure S3**.

### Quantitative proteomics analysis of SASP composition

To gain a comprehensive profile of the heterogenous SASP composition and maximize the detection of SASP factors important for reprogramming other than IL6, we isolated the soluble protein fractions from the OIS-CM and Ctrl-CM of *Il6^-/-^* MEFs to perform quantitative proteomic analysis. The label-free data-independent acquisition (DIA) tandem mass spectrometry strategy allows sensitive and unbiased detection of SASP proteins in a reproducible manner (Basisty et al., 2020; Gillet et al., 2012). To ensure the correct identification, we only included a protein when it can be detected in all six replicates of one condition. The proteomic analysis identified a total of 1955 proteins in both conditions, including 249 proteins with a significant differential expression (q-value <0.01) that had a fold change of at least 2-fold (SEN/CTL) (Figure 3B). Of note, 217 proteins were exclusively detected in one condition (185 in SEN and 32 in CTL) (Figure 3C), which were assumed differentially abundant between the conditions and included in the data analysis. We found many known SASP factors are more abundantly secreted in the OIS-SN, such as SPP1, PLAU, and MMP3 (Figure S3D). Of note, we did not detect IL-6 in both conditions. Interestingly, there is less than 50% overlap between our study and a recent large-scale human SASP proteomic study (human fibroblast-OIS, 115/386), suggesting heterogenous in SASP composition between species (Hernandez-Segura et al., 2017).

To identify which biological pathways/responses are mediated by SASP factors, we performed pathway and network analyses on the proteins that are either more abundantly or exclusively secreted from the SEN condition. We found the most dominant clusters of gene ontology (GO) pathways are related to morphogenesis and stress response, including cell morphogenesis in development and differentiation, response to growth factor, wounding response, and angiogenesis (Figure 3D). Besides, the hallmark gene set enrichment analysis (GSEA) revealed several pathways that are associated with paracrine senescence, including Epithelial-to-Mesenchymal transition (EMT) (Ansieau et al., 2008), TNFα signaling via NFκB, inflammatory response, and p53 pathway (Coppe et al., 2010) (Figure S3E).

### Amphiregulin enhances *in vitro* reprogramming efficiency and kinetic

Of note, several pathways identified by the bioinformatic analysis are also important for somatic reprogramming, such as EMT (Liu et al., 2013), apoptosis (Kawamura et al., 2009; Kim et al., 2018), hypoxia (Yoshida et al., 2009), and p53 pathway (Hong et al., 2009; Kawamura et al., 2009; Marion et al., 2009; Utikal et al., 2009). Given Mesenchymal-to-Epithelial transition (MET) is required in the initiation phase of *in vitro* reprogramming (Li et al., 2010; Liu et al., 2013; Samavarchi-Tehrani et al., 2010), and because our data suggested that paracrine senescence promotes reprogramming at the early stage (Figure 1D), we investigated the proteins overlapped with the EMT gene set and found many factors crucial for *in vitro* reprogramming, such as BMP1 and TGFβ (Li et al., 2010; Liu et al., 2013; Samavarchi-Tehrani et al., 2010)(Figure S4A).

AREG is an EGFR ligand (Shoyab et al., 1989) and has diverse functions in development, tissue regeneration, and tumorigenesis (Zaiss et al., 2015). Since both OIS and muscle injury strongly induce AREG (Burzyn et al., 2013; Pommer et al., 2021), we wondered whether AREG could partially mediate the effect of paracrine senescence on reprogramming both *in vitro* and *in vivo*. Interestingly, AREG is the only EGFR ligand that has significantly increased mRNA expression in OIS comparing to IRIS (Figure S4B). To formally test the effect of AREG on reprogramming, we added different concentrations of the recombinant AREG protein for the first three days during reprogramming, consistent with the CM experiment (Figure 1D). AREG treatment enhanced reprogramming efficiency in a dosage-dependent manner (Figure 4A). We next removed DOX at different time points to examine the ability of AREG to facilitate OSKM transgene-independent clonal growth, a hallmark of iPSCs. AREG treatment yielded iPSCs clones after only 5 days of OSKM expression, compared to 8 days in the control condition (Figure 4B), suggesting a faster reprogramming kinetic.

**Figure 4.**
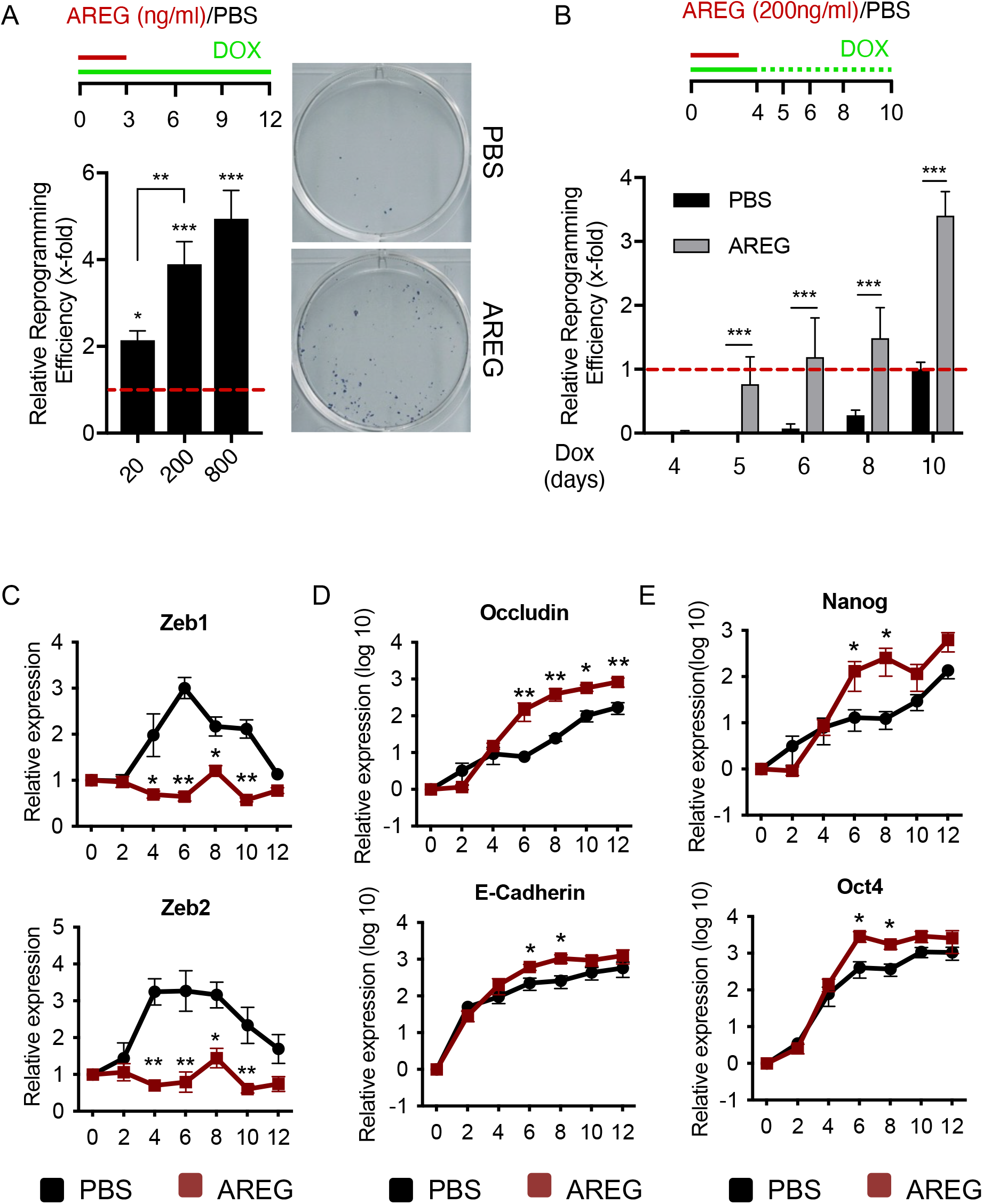
AREG promotes *in vitro* reprogramming. **A.** Experimental scheme (upper panel). Fold change of reprogramming efficiency of i4F MEFs with different dosages of recombinant AREG protein. Normalized to Vehicle control (PBS). Representative image of plate stained with alkaline phosphatase (AP) to reveal colonies arising from reprogramming (right panel). **B.** Experimental scheme (upper panel). Comparison of reprogramming efficiency between PBS and recombinant AREG after different duration of DOX treatment. Data correspond to the average ± s.d. using indicated n number of independent experiments. **C-E.** Kinetics of expression of indicated genes at the indicated days during reprogramming comparing between PBS and AREG as measured by quantitative RT-PCR. Data correspond to the average ± s.e.m. of at least 2 independent assays. For each assay, qRT-PCR values were obtained in duplicate. Statistical significance was assessed by ordinary one-way ANOVA test for **A**, and two-way ANOVA test for **B-E:** *p < 0.05; **p < 0.01; ***p < 0.001. See also **Figure S4**.

### AREG accelerates cell cycle and the process of MET during reprogramming

We sought to understand the mechanism by which AREG promotes reprogramming. The reported role of AREG in proliferation and EMT (Berasain and Avila, 2014) prompted us to analyze cell proliferation and MET process, two vital cellular events that occurred in the early stage of reprogramming (Sancho-Martinez and Izpisua Belmonte, 2013). We found that AREG treatment led to a larger fraction of cells in S-phase and increased cell proliferation globally in MEFs alone (Figure S4C & S4D). Next, we checked whether AREG affected MET during reprogramming by measuring the expression of several mesenchymal and epithelial cell markers at different time points throughout reprogramming using qPCR. In control, the expression of the mesenchymal markers, Zeb1 and Zeb2, were transiently upregulated and gradually suppressed afterward. Interestingly, these markers were suppressed throughout the reprogramming process in AREG treated condition (Figure 4C). Conversely, the expression of the epithelial markers, Occludin and E-Cadherin, was higher in samples treated with AREG than the control (Figure 4D). Besides, the expression of all the pluripotency markers, endogenous Oct4, Sox2, Klf4, Nanog, Lin28, and Esrrb, gradually increased and peaked at the last stage reprogramming in control. While in AREG treated condition, endogenous Oct4 and Nanog expression sharply increased at 6 days and reached the plateau earlier than in the control (Figure 4E). However, the expression dynamic of other pluripotent markers remained the same (Figure S4D), suggesting an action upon Nanog and Oct4 specifically rather than a mere reflection of the accelerated reprogramming. Taken together, these results indicate that AREG enhances the efficiency and kinetic of reprogramming by increasing proliferation rates and facilitating MET process.

### EGFR signaling pathway is important for reprogramming

Since AREG is an atypical EGFR ligand (Zaiss et al., 2015), we wondered whether other EGFR ligands could also promote somatic reprogramming and found that additional epidermal growth factor (EGF) also improved the reprogramming efficiency (Figure 5A). Next, we examined whether EGFR activation is necessary for the enhancement. AREG and Dox combination strongly induced the activation of EGFR comparing to Dox alone or untreated control determined by the level of the phosphorylated EGFR using Western blotting (Figure S5A). Besides, Lapatinib, a dual tyrosine kinase inhibitor of EGFR and human epidermal growth factor receptor-2 (HER2) (Wood et al., 2004), canceled the effect of AREG on reprogramming (Figure 5B and Figure S5A). Of note, a high concentration of Lapatinib alone significantly repressed reprogramming (Figure S5B), suggesting EGFR signaling pathway might be required for reprogramming. Hence, AREG promotes reprogramming in an EGFR dependent manner.

**Figure 5.**
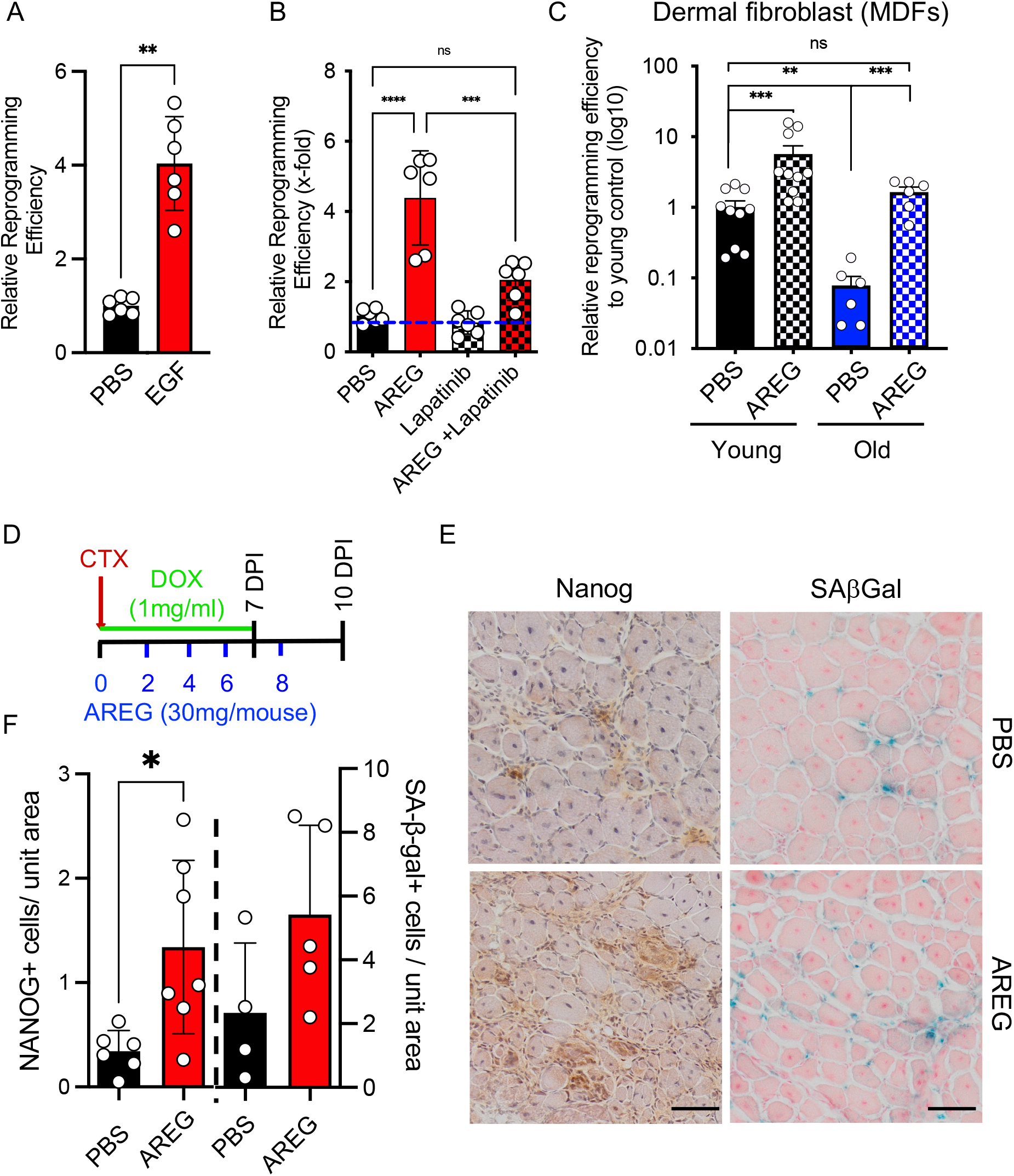
AREG promotes *in vivo* reprogramming. **A.** Relative reprogramming efficiency of i4F MEFs treated with EGF (100ng/mL). Relative to PBS treated sample. **B.** Relative reprogramming efficiency of i4F MEFs with indicated treatment. Relative to PBS treated sample. Data correspond to the average ± s.d. of using indicated n number of different experiments using i4F MEFs generated from different embryos (A & B). **C.** Reprogramming efficiency of dermal fibroblasts derived from i4F mice with the indicated treatment and age. Data correspond to the average ± s.d. of using indicated n number of different MDF isolates. **D.** Experimental scheme for AREG administration during *in vivo* reprogramming. **E.** Representative TA sections with immunohistochemistry staining of anti-Nanog and histological staining of SAβGal of indicated treatment. Scale bar: 50 μm. **F.** Quantification of E. Data correspond to the average ± s.d. of using indicated n number of different mouse (average of two TA per mouse). Statistical significance was assessed by the Mann-Whitney test (A) and ordinary one-way ANOVA (B, C, and F): *p < 0.05; **p < 0.01; ***p < 0.001. See also **Figure S5**.

### AREG rescues age-dependent reduction in reprogramming

Previously, we reported that murine dermal fibroblasts (MDFs) derived from old (> 2 years) mice had a significant reduction in reprogramming efficiency comparing to MDFs derived from young (2 months) mice (Li et al., 2009). Because AREG induces cell proliferation, we investigated whether AREG can improve reprogramming in this context. Consistent with the previous report, MDFs from old (> 2 years) i4F mice had much lower reprogramming efficiency comparing to MDFs from young (2 months) i4F mice (Figure 5C). Notably, the addition of AREG to the aged MDFs rescued their low reprogramming efficiency to the same level as the young MDFs (Figure 5C).

### AREG promotes *in vivo* reprogramming in the skeletal muscle

It has been reported that AREG enhances the myogenic differentiation of skeletal muscle stem cells (MuSCs) *in vitro* and facilitates muscle regeneration (Burzyn et al., 2013). We previously reported that MuSC is a cell of origin for *in vivo* reprogramming (Chiche et al., 2017). Therefore, we hypothesize that AREG might also improve *in vivo* reprogramming in the injured muscle. To address this question, we administered AREG to the CTX-injured, DOX-treated male i4F mice according to the experimental scheme illustrated in Figure 5D and evaluated reprogramming and senescence induction at 10 DPI. (Note that the initial AREG injection was done i.m. as previously described (Burzyn et al., 2013).) There were many more Nanog+ cells in the TA sections from mice treated with AREG comparing to the control group (Figure 5E and Figure 5F). Conversely, there is no apparent difference in senescence induction, detected by SAβGal staining (Figure 5E and Figure 5F). Because AREG was administrated mainly i.p., we wondered whether it could also improve reprogramming in other tissues. Interestingly, we also observed more Nanog+ cells in both pancreas and kidney, but not in the liver (Figure S5C), suggesting a tissue-specific effect of AREG on *in vivo* reprogramming.

## Discussion

Here, we report that the impact of paracrine senescence on reprogramming is stress-dependent. The quantitative proteomic analysis on the senescence-associated secretome identified a catalog of factors involved in various biological processes that could potentially impact cell plasticity. We demonstrated that the AREG enhances *in vitro* reprogramming efficiency and kinetic, at least in part, by accelerating somatic cell proliferation and the process of MET. Of note, AREG treatment rescue the age-associated reduction in reprogramming and facilitate *in vivo* reprogramming in a tissue-specific manner.

IL-6 is required for oncogene-induced senescence (Kuilman et al., 2008). Surprisingly, we could efficiently induce senescence in *Il6^-/-^* MEFs (Figure S2A)(Kopf et al., 1994). Both ELISA and proteomic analysis failed to detect IL-6 in the CM-OIS-*Il6^-/-^* (Figure S2B and Figure S2C), indicating that OIS-*Il6^-/-^* does not secrete IL-6. Thus, the discrepancy might be due to the compensation effect of germline knockout, which often yields much weaker phenotypes than acute disruption of gene function (Housden et al., 2017). Importantly, IL-6 blocking antibody did not abolish the beneficial effect of CM-OIS-*Il6^-/-^* on *in vitro* reprogramming (Figure 2B), highlighting the involvement of other SASP factors.

Consistently, we found that IL-6 deficiency supports *in vivo* reprogramming in the muscle (Figure 2C and Figure 2D). Together with others, we have previously demonstrated that tissue injury-induced senescence facilitates *in vivo* reprogramming in a paracrine manner, particularly by IL-6 (Chiche et al., 2017; Mosteiro et al., 2016). A recent study reported a reduction of reprogramming in the pancreas of i4F; *Il6^-/-^* mice (Mosteiro et al., 2018). Firstly, reprogramming-induced and injury-induced senescence might produce different factors (Chiche et al., 2017; Mosteiro et al., 2018). Besides, IL-6 is a myokine vital for muscle formation and growth (Serrano et al., 2008). Although many different cell types produce IL-6 upon muscle injury, including macrophages and muscle stem cells (Munoz-Canoves et al., 2013), muscle regeneration is only mildly affected in *Il6^-/-^* mice (Zhang et al., 2013), suggesting a potential compensation by other members of the IL-6 cytokine family. Therefore, the tissue-specific requirement of IL-6 for *in vivo* reprogramming might be due to distinct stress/tissue-dependent senescence secretome and the pleiotropic functions of IL-6.

To gain an extensive and quantitative view on SASP factors, we performed an unbiased proteomic study using CM-OIS-*Il6^-/-^* (Figure 3B). The bioinformatic network analysis revealed pathways predominantly related to morphogenesis and response to growth factors (Figure 3C), where EGFR signaling pathways play crucial roles. Of note, we used OIS-MEFs as a proxy for senescence induced by OSKM over-expression or muscle injury to identify SASP factors, which might not fully recapitulate the *in vivo* scenario. Future investigation will further elucidate the context-dependent SASP heterogeneity.

To shed some light on SASP factors other than IL-6 and reprogramming, we investigated the role of AREG in this process. We found that AREG treatment could notably increase *in vitro* reprogramming efficiency (Figure 4A) and kinetic (Figure 4B). Next, we showed that AREG could accelerate cell proliferation (Figure S4C) and suppress mesenchymal markers (Figure 4C), suggesting AREG might affect several cellular events necessary for the initiation phase of reprogramming(Sancho-Martinez and Izpisua Belmonte, 2013). Interestingly, we found AREG could alleviate the negative impact of age on reprogramming (Figure 5B), providing a straightforward solution to improve iPS cell generation during aging (Mahmoudi et al., 2019). Of note, AREG treatment failed to further enhance reprogramming to the similar level as the young (Figure 5B), reflecting the attenuated EGFR signaling in the aged skin fibroblasts as previously reported (Shiraha et al., 2000; Tran et al., 2003).

Furthermore, we showed that AREG promotes *in vivo* reprogramming in the muscle (Figure 5E & 5F). It has been reported that EGFR regulates the asymmetrical division of MuSCs (Wang et al., 2019), and AREG could facilitate the differentiation of MuSCs *in vitro* (Burzyn et al., 2013; Zaiss et al., 2015). Therefore, AREG might directly promote the cell plasticity of MuSCs, a primary cell origin of reprogramming in the muscle (Chiche et al., 2017). Interestingly, AREG also promoted reprogramming in the pancreas and kidney but not in the liver (Figure S5C), highlighting a tissue-specific mode of action. Of note, recent studies showed that cyclic expression of OSKM could enhance muscle regeneration (Ocampo et al., 2016). Given senescence is important for optimal wound healing (Demaria et al., 2014) and limb regeneration (Yun et al., 2015), future investigation is warranted to determine whether the rejuvenation effect of partial reprogramming is partially mediated by senescence.

In summary, we dissected the impact of paracrine senescence on reprogramming and revealed the role of the AREG-EGFR signaling pathway in reprogramming. These findings may have important implications for understanding senescence’s involvement in tissue repair and *in vivo* reprogramming.

## Acknowledgements

We are indebted to Jun Zhang for his excellent technical support in image analysis. We are grateful to the Central Animal Facility, the Proteomic Platform, the Cytometry Platform, and the Bioinformatics and Biostatistics Hub of the Institut Pasteur. Work in the laboratory of H.L. was funded by Institut Pasteur, Centre National pour la Recherche Scientific and the Agence Nationale de la Recherche (Laboratoire d’Excellence Revive, Investissement d’Avenir; ANR-10-LABX-73; ANR-16-CE13-0017). This work is also supported by Fondation ARC (PJA 20161205028). AC was funded by the postdoctoral fellowships from the Revive Consortium.

## AUTHOR CONTRIBUTIONS

M.VJ performed most of the experimental work, contributed to experimental design, data analysis, and discussions. C.C made critical experimental contributions. AC contributed experimentally. T.D performed proteomic analysis and data acquisition. M.M supervised the proteomic analysis and data interpretation. H.L. supervised the study, designed the experimental plan, interpreted the data and wrote the manuscript. All authors discussed the results and commented on the manuscript. The authors declare no competing financial interests with this paper.

