## Supplementary Information for "Amphiregulin mediates non-cell-autonomous effect of senescence on reprogramming" (from another version of this preprint: version 1, 2021-09-01)

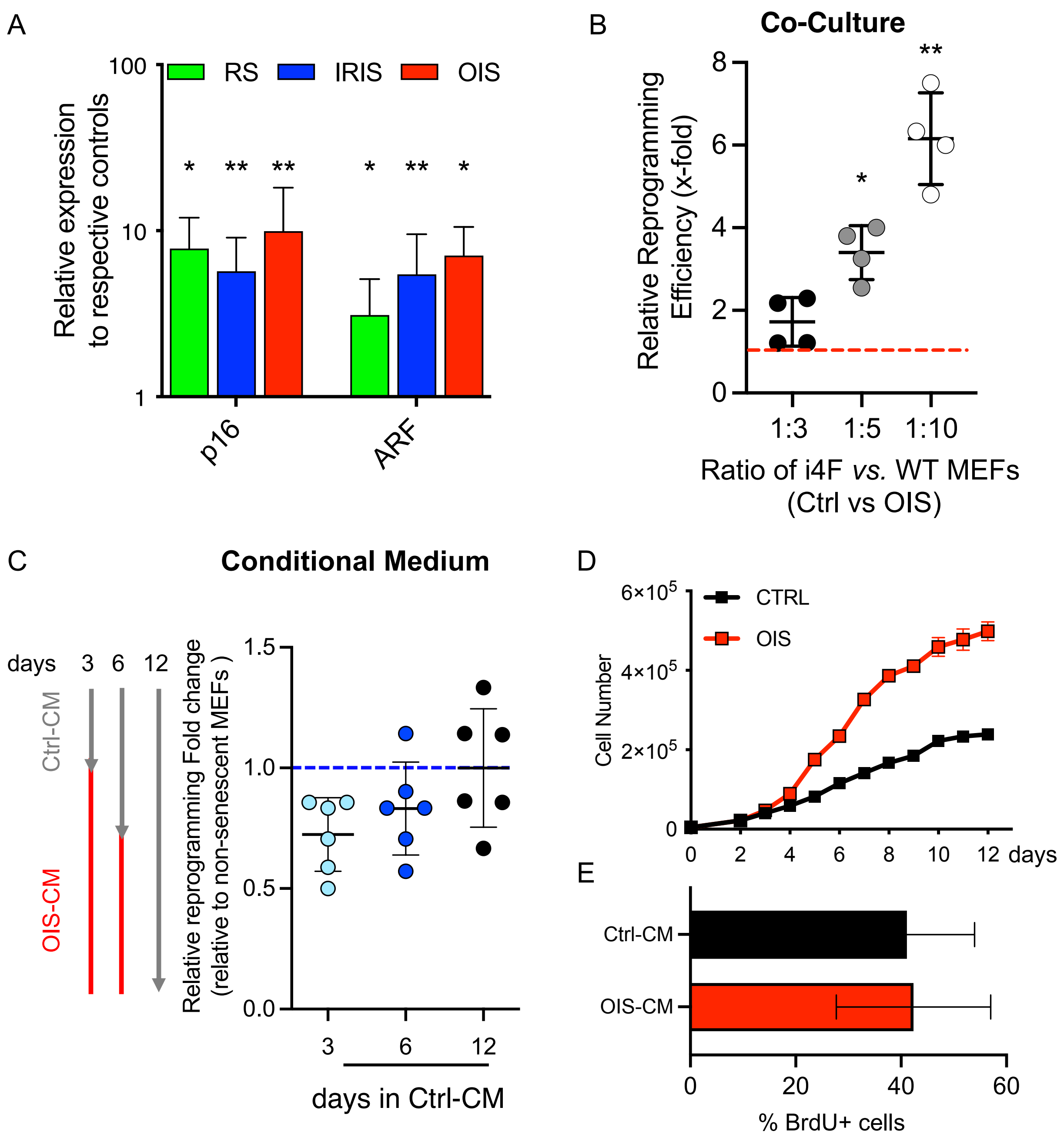


***Supplemental Figure 1 Related to Figure 1***

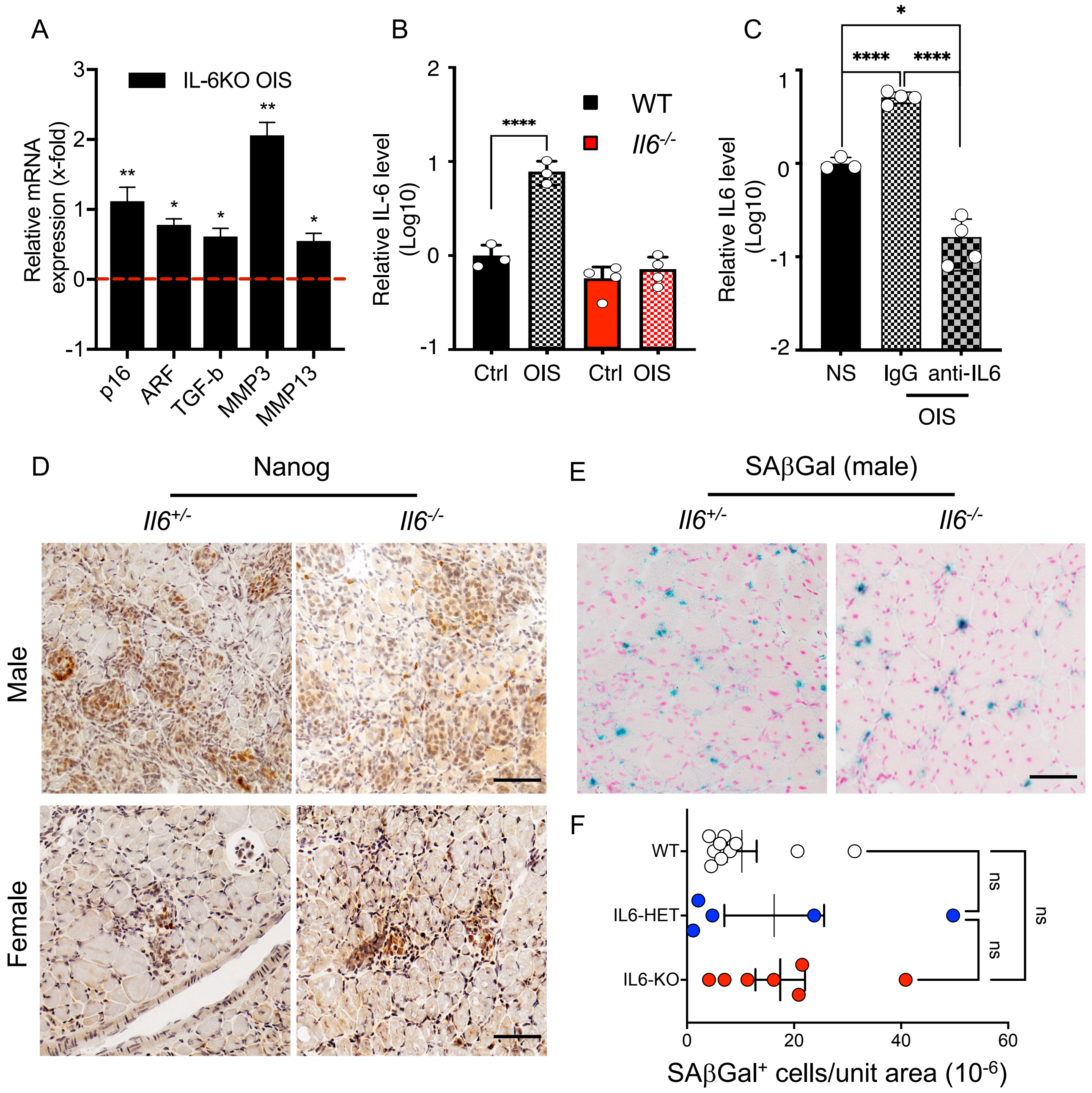


***Supplemental Figure 2 Related to Figure 2***

A B C

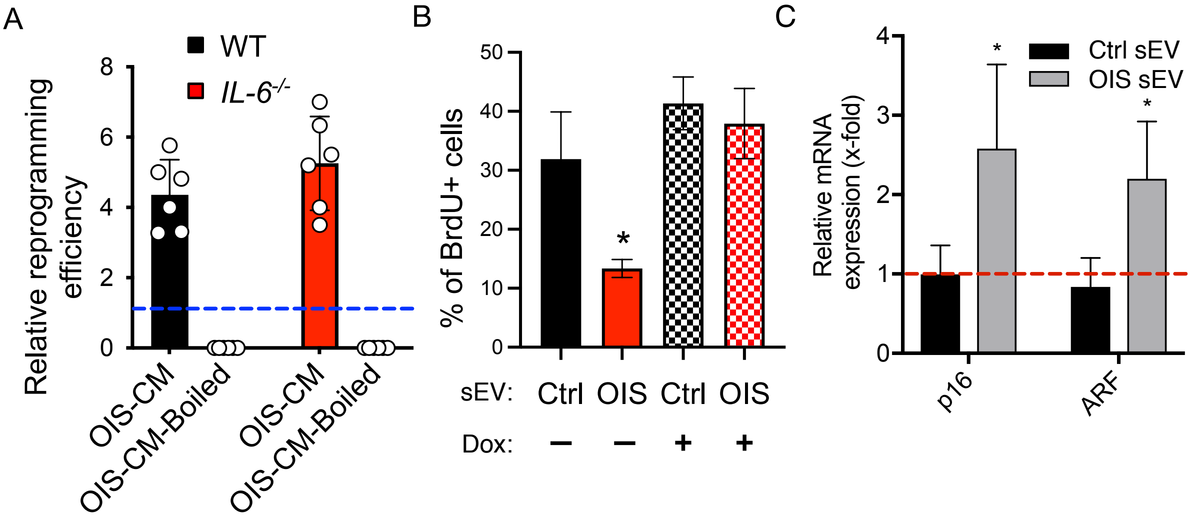


D Top 10 upregulated SASP (OIS> Ctrl)

**Bold:** absent in the SASP atlas (www.saspatlas.com)

| Names | LogFC (OIS/Ctrl) |
|---|---|
| **SPP1** | 9.92 |
| PLAU | 9.73 |
| **NPTX1** | 8.07 |
| MMP3 | 6.68 |
| **PLA2G7** | 6.27 |
| GPC1 | 5.75 |
| **CRABP1** | 5.51 |
| **TSLP** | 5.45 |
| **SEMA4B** | 5.18 |
| HSP90B1 | 5.12 |

E Top 20 enriched pathways (Hallmark gene sets)

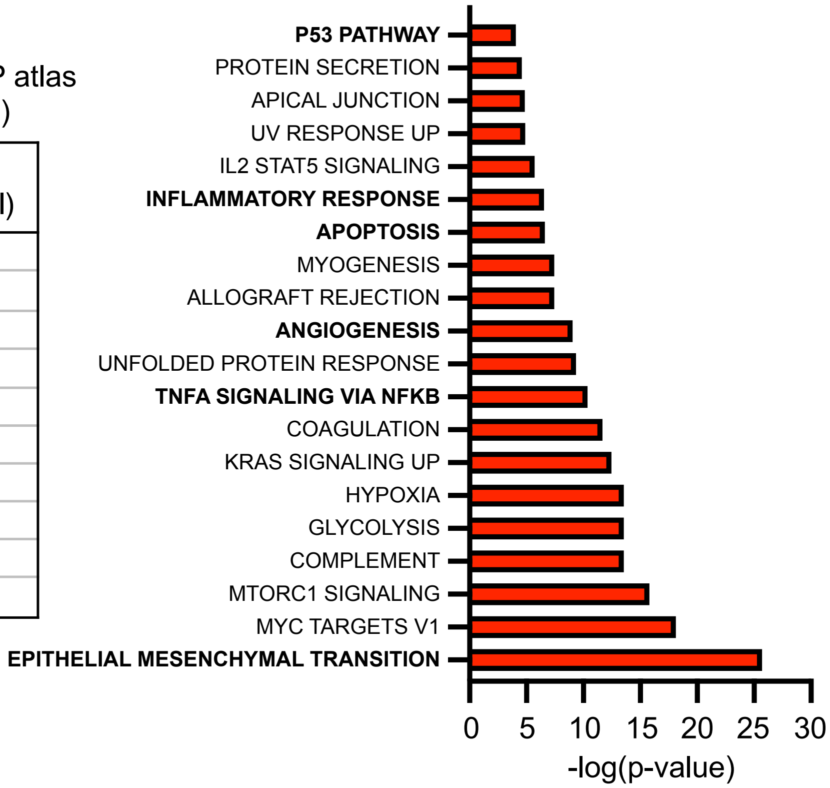


***Supplemental Figure 3 Related to Figure 3***

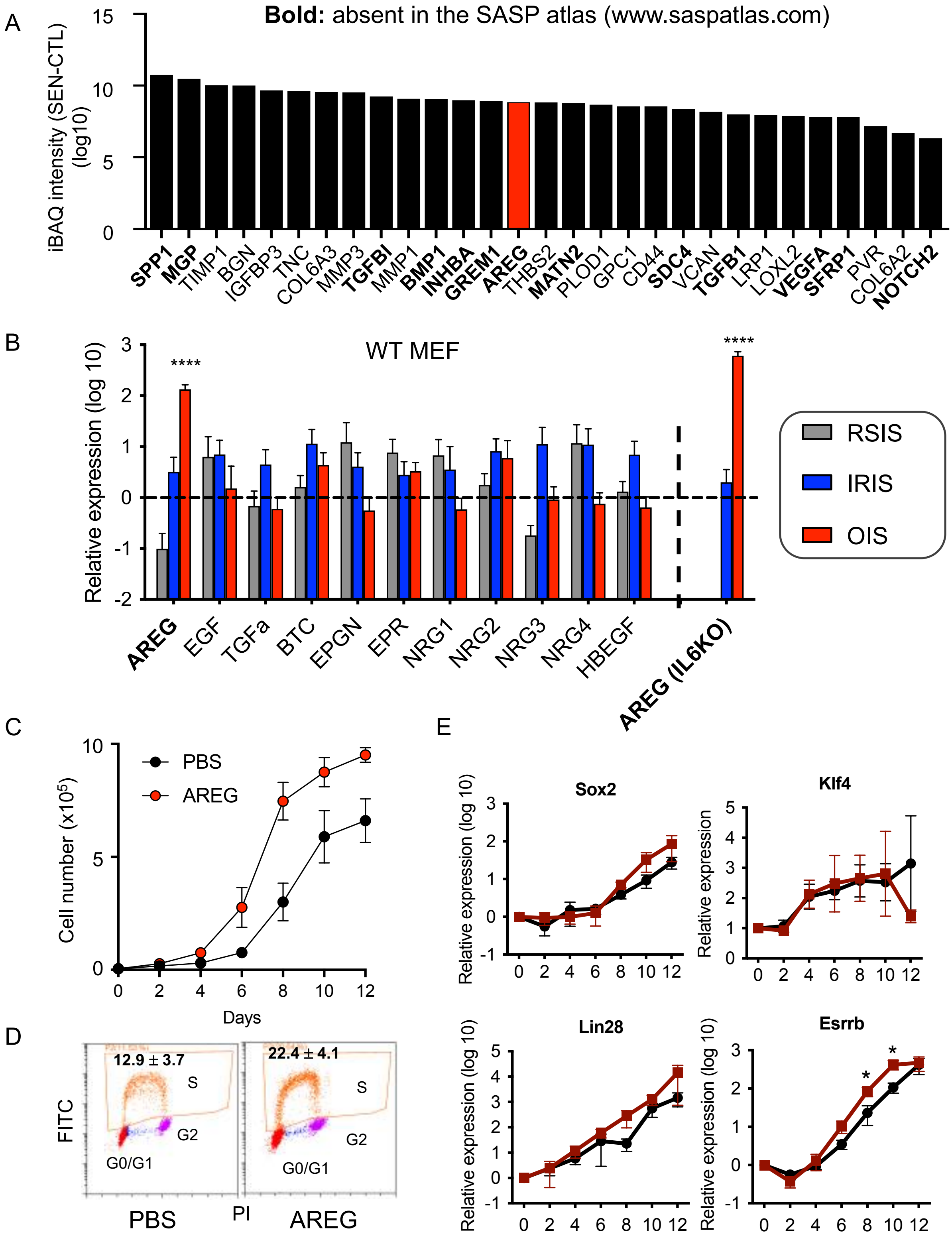


**Supplemental Figure 4 Related to Figure 4**

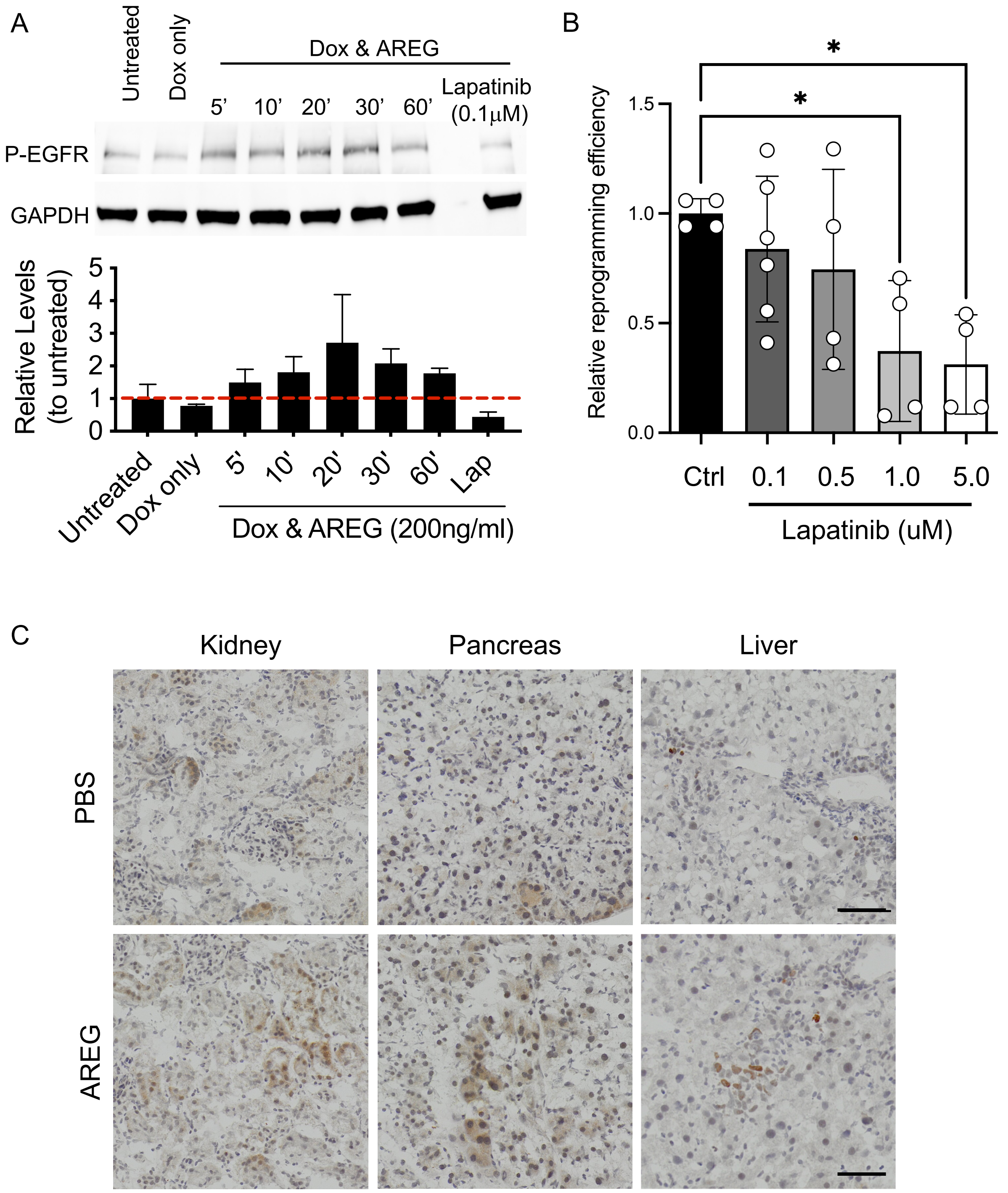


***Supplemental Figure 5 Related to Figure 5***

### SUPPLEMENTAL FIGURE LEGENDS

**Figure S1. Related to Figure 1. Senescence promotes *in vitro* reprogramming in a stress-dependent manner**

**A.** Expression of *p16* and *Arf* in different stress-induced senescent WT MEF compared with the non-senescent experimental control measured by quantitative RT-PCR (qRT-PCR). **B.** Fold change in the reprogramming efficiency of i4F MEFs co-cultured with either control or OIS treated WT MEFs. Different amount of i4F MEFs were seeded on top of $2x10^5$ cells with ratios of 1:3, 1:5, and 1:10 (i4F: WT). **C.** Reprogramming efficiency of i4F MEFs cultured with different duration of Ctrl-CM before switching to OIS-CM. Experimental scheme (left panel). All the reprogramming fold changes was normalized to i4F in Ctrl-CM for 12 days, indicated by the blue dash line. Data correspond to the average ± s.d. using indicated n number of independent experiments. **D.** Growth curve of WT MEFs cultured in either Ctrl-CM or OIS-CM for 12 days. Cell numbers were counted every two days. Data correspond to two independent experiments with triplicates of each condition per experiment. **E.** Quantification of the percentage of BrdU+ cells after 72 hours incubation with either Ctrl or OIS-CM. Data correspond to five immunofluorescence images per condition. Statistical significance was assessed by ordinary by the Mann-Whitney test and one-way ANOVA test: *$p < 0.05$; **$p < 0.01$.

**Figure S2. Related to Figure 2. *in vivo* reprogramming in the skeletal muscle occurs normally in *Il6*$^{-/-}$ mice**

**A.** Expression of senescence-associated genes in *Il6*$^{-/-}$ MEFs upon oncogene-induced senescence compared with the non-senescent experimental control measured by qRT-PCR. **B & C.** Relative fold change in IL-6 protein level measured by ELISA in indicated genotypes and conditions. WT compared to *Il6*$^{-/-}$ MEFs (**B.**) and IL-6 blocking antibody (**C.**) Data correspond to the average ± s.d. of using indicated n number of experiments. **D & E.** Representative TA sections with immunohistochemistry staining of anti-Nanog (**D**) and histological staining of SAβGal (**E**) of indicated genotypes and sex. Scale bar: 50 μm. **F.** Quantification of E. Data correspond to the average ± s.d. of using indicated n number of different mouse (average of two TA per mouse). Statistical significance was assessed by the ordinary one-way ANOVA: *$p < 0.05$; **$p < 0.01$; ****$p < 0.0001$.

**Figure S3. Related to Figure 3. Proteomic analysis of oncogenic stress-induced secretome in *Il6*$^{-/-}$ MEFs**

**A.** Fold change in reprogramming efficiency of i4F MEFs cultured in either complete CM or boiled CM of indicated genotypes. Efficiency was normalized to CM-Ctrl of respective genotype and indicated with the blue dash line. Data correspond to the average ± s.d. using indicated n number of independent experiments. **B.** The percentage of BrdU+ cells of i4F MEFs treated with various indicated conditions. sEV: small extracellular vesicles (exosome). Dox: Doxycycline. Data correspond to the average quantification of the immunofluorescence analysis of BrdU staining of two independent experiments. **C.** Expression of *p16* and *Arf* in WT MEFs treated with either non-senescent exosome (Ctrl-sEV) or OIS exosome (OIS-sEV). The expression was compared to Ctrl-CM and measured by qRT-PCR. **D.** List of top 10 most upregulated secreted proteins in OIS comparing to Ctrl identified by the proteomic analysis. **E.** Hallmark gene set enrichment analysis shows various pathways overrepresented among proteins specifically or differentially released by OIS- *Il6*$^{-/-}$ MEFs. Statistical significance was assessed by ordinary by the Mann-Whitney test (A, B & C) and one-way ANOVA test: *$p < 0.05$.

**Figure S4. Related to Figure 4. AREG promotes in vitro reprogramming**

**A.** Increase in protein intensity in OIS-CM compared to Ctrl-CM of the proteins presented in the Epithelial-Mesenchymal-Transition pathway. Protein intensity was measured by the total intensities divided by the identified peptides for one protein (iBAQ). **B.** Expression of EGF-family ligands in different stress-induced senescent WT MEF compared with the non-senescent experimental control measured by quantitative RT-PCR. The expression of AREG was also measured senescent *Il6*$^{-/-}$ MEFs by qRT-PCR. **C.** Proliferation curve of i4F MEFs cultured in either PBS or AREG (200ng/ml, treated for the first three days) in absence of DOX. **D.** Cell-cycle distribution analyzed by BrdU incorporation of the propidium iodide (PI)-positive and -negative populations of cells obtained 3 days after AREG (200ng/ml) treatment or PBS (control). Two independent experiments were performed, and 10000 cells are represented in each dot plot. **E.** Kinetics of expression of indicated genes at the indicated days during reprogramming comparing between PBS and AREG as measured by quantitative RT-PCR. Data correspond to the average ± s.e.m. of at least 2 independent assays. For each assay, qRT-PCR values were obtained in duplicate. Statistical significance was assessed by ordinary by the Mann-Whitney test (B & E) and one-way ANOVA test: *p < 0.05; **p < 0.01; ****p < 0.0001.

**Figure S5. Related to Figure 5. AREG promotes in vivo reprogramming**
**A.** Western-blot analysis of EGFR activation upon different treatments indicated. The quantification of the phosphorylation form of EGFR normalized to the GAPDH loading control is showed in the lower panel. **B.** Reprogramming efficiency of i4F MEFs treated with various concentration of Lapatinib (µM). **C.** Representative tissue sections with immunohistochemistry staining of anti-Nanog of indicated tissue and condition. Scale bar: 50µM.

### Supplemental Experimental Procedures:

**Mice and breeding.** Animals were handled as per European Community guidelines and the ethics committee of the Institut Pasteur (CETEA) approved protocols. The reprogrammable i4F mice were kindly provided by Manuel Serrano (Spanish National Cancer Research Centre, Madrid, Spain) (Abad et al., 2013). The $Il6^{-/-}$ were described previously (Kopf et al., 1994) and were crossed with i4F. To induce muscle injury, mice were anesthetized with isoflurane. *Tibialis anterior* (TA) muscles were injured by injection 40µl of snake venom cardiotoxin (10µM) (L8102, Lotaxan Valence, France, www.latoxan.com). *In vivo* reprogramming was induced by administration of Doxycycline (Sigma) in the drinking water supplemented with 7.5% of sucrose right after injury. Experiments were performed indistinguishably with male mice from 6 to 8 weeks of age except indicated otherwise.

**Animal procedures.** For AREG antibody injection, after muscle injury by cardiotoxin, mice were intraperitoneally injected by antibodies against AREG (989-AR, R&D systems) or purified Rat IgG1 control (BioLegend, clone RTK2071) at the concentration of 7 $\mu$g/mouse at days 0, 2, 4, 6 and 8-post injury and TAs were collects at 10 days-post injury. The initial AREG injection was done intra-muscularly.

**Cell culture conditions**
Primary mouse embryo fibroblasts (MEFs, passage 2) were obtained from pure inbred C57BL6 background WT or reprogrammable mice, as described previously (Li et al., 2009). Mouse dermal fibroblasts (MDFs) were obtained from the ear of young (2 months) and old (2 years) i4F mice as described before (Li et al., 2009). Fibroblasts (MEFs and MDFs) were cultured in standard DMEM medium with 10% FBS (Gibco) and 1% Pen/Strep (Gibco).

**Induction of senescence.** γ-irradiation induced senescence (IRIS): WT MEFs were treated with ionizing radiation (10 Gy X-ray). Quiescent control cells were mock irradiated. Oncogene Induced Senescence (OIS): MEFs were infected with hRAS viral particles or control vector particles. Following infection, cells were selected with puromycin (gene resistance marker) for 4 days and seeded at proper density on tissue culture plates previously treated with gelatin 0,1% (10 min at RT) for further experiments. Replicative Senescence (RS): MEFs were propagated every 3 days under 3% $O_2$ and 5% $CO_2$ (hypoxia conditions). Cells were seeded at 20% confluency and passed upon reaching 80-90% confluency. This protocol was performed until cells reached replicative crisis and they stopped growing. Then cells were seeded at proper density on tissue culture plates previously treated with gelatin 0,1% for further experiments. Media was changed every 2 days. Senescent cells were cultured for 10 days to allow development of the senescent phenotype, and quiescent cells were cultured in 0.2% serum for 3 days.

**Collection of CM.** Senescent cells were then washed with PBS (Thermo Fisher Scientific) and placed in serum- and phenol red–free DMEM (Thermo Fisher Scientific, Waltham, MA; #21063–029), and conditioned media was collected after 48 hours and filtered through 0,2µm filter.

**Soluble fraction and sEV isolation.** Whole CM was centrifuged at 500g for 10min filtered through 0.2µm filter unit. Next, filtered CM was centrifuged at 12 000g for 30 min at 4°C, pellet was removed, and supernatant was centrifuged at 100 000g for 3 hours to collect the soluble fraction (SN) absence of exosome. The pellet was resuspended with PBS and centrifuged at 100 000g for additional 3 hours. The purified pellets were collected and resuspended into proper medium for further experiments without further concentration. Samples were centrifuged using an Ultracentrifuge Optima XPN-80 Ultracentrifuge from Beckman Coulter equipped with a rotor SW32Ti.

***In vitro* reprogramming in co-culture & CM system.**
For the co-culture system, primary i4F MEFs were seeded over senescent or quiescent cells described above at density of $2x10^4$ per well in 6-well plate. For conditioned medium system, primary i4F MEFs were seeded at the density of $5x10^3$ per well in 6-well plate. *In vitro* reprogramming was induced on the second day by changing to the iPSCs medium of high-glucose DMEM supplemented with KSR (15%, Invitrogen), LIF (1,000 U/ml), non-essential amino acids, penicillin-streptomycin, glutamax and -mercaptoethanol) supplement with doxycycline (1 µg/ml) as described previously (Abad et al., 2013). For the CM system, conditioned medium from either senescent or quiescent cells were used instead of high-glucose DMEM and the rest components were added as described above. Medium was changed every 48h until iPS cell colonies appeared. For the IL-6 blocking antibody experiment, IgG antibody (1µg/mL, BioLegend, purified Rat IgG1, κ Isotype Ctrl Antibody) and anti-IL-6 antibody (3µg/mL, eBioscience clone MP5-20F3) were used. For AREG, EGF, and Lapatinib experiment, AREG (200ng/ml excepted indicated otherwise, R&D Systems, 989-AR), EGF (R&D Systems, 2028-EG), and Lapatinib (Tebu-Bio, T0078) were used for the first three days. Medium were changed every 24h until iPS colonies appeared. PBS were used as the control. Reprogramming plates were stained for alkaline phosphatase activity (Sigma, AP detection kit AB0300) and quantified using ImageJ software.
For the proliferation analysis, $2x10^3$ cells of primary i4F MEFs were seeded in triplicates onto wells of 12-well plates. Cell number were counted using Neubauer chamber every two days for 10-12 days period. Medium were changed every 24h.

**Plasmids and infections.** Retroviral vectors pbabe-hRASV12-puro (Addgene #9051) and pbabe-puro (Addgene #1764) (control) were used for infections of MEFs. HEK 293 cells were transfected using PEI (Polyethylenimine) with a 1:4 ratio of DNA:PEI. MEFs were infected twice per day for 2 days (12h interval). Viral supernatant was collected, centrifuged at 300g, and filtered through 0,45µm. Polybrene was used (8µg/mL) (Sigma, TR-1003). Following day after infection, cells were selected with puromycin (4µg/mL) (Sigma, P8833).

---

**Proteomic sample preparation**
**Chemicals.** Acetonitrile (CAN, #34851), water (#270733), Iodoacetamide ((#I114), Tris(2-carboxyethyl)phosphine hydrochloride (TCEP, #646547), Trichloroacetic acid (TCA, #T6399), ammonium bicarbonate (#09830) were from Sigma-Aldrich (USA), formic acid (#94318, Fluka), Urea (#29700) was from Thermo Scientific (Waltham, MA), sequencing grade trypsin (#V5113) was from Promega (San Luis Obispo, CA), and Sep-Pak C18 cartridge (50mg sorbent, #WAT054955) were from Waters (Milford, USA).

**Sample collection.** OIS and mock-infected cells were washed with DMEM twice before incubation with DMEM (without FBS and Pen/Strep) for 48 hours at 37°C under normal conditions. CM was collected, centrifuged 10 min at 500g, filtered through 0,2µm filter unit and centrifuged at 12 000g for 30 min at 4°C. Pellets (containing sEVs) were removed. CM was centrifuged for additional 3 hours at 100 000g. CM was kept frozen for further analysis.

**Digestions.** Two technical replicates were processed for every sample. Proteins from the secretome were precipitated 1hour at 4°C with TCA (20%) and centrifugated 15min at 16 000g in 4°C. The pellets were washed twice with ice-cold acetone and resuspended in denaturation buffer containing 8M Urea/100mM ammonium bicarbonate. The mixtures were reduced with 50mM TCEP (37°C for 1 hour), then alkylated with 50 mM iodoacetamide (1 hour at RT in the dark). Samples were diluted 10-fold with 100mM ammonium bicarbonate and incubated overnight at 37°C with sequencing grade trypsin (Promega, San Luis Obispo, CA) at a 1:40 enzyme:substrate ratio (wt/wt).

**Desalting.** 4% FA was added to stop the reaction. Digested peptides were purified with Sep-Pak C18 and were eluted with 80% (ACN) containing 0.1% formic acid in water. Resulting peptides were dried and resuspended in 2%ACN containing 0.1% formic acid in water.

**Mass spectrometry analysis.** Samples acquisitions were performed on a Q Exactive™ Plus Mass Spectrometer (Thermo Fisher Scientific, USA) coupled with a Proxeon EASY-nLC 1200 (Thermo Fisher Scientific, USA). 1$\mu$g of peptides were injected onto a home-made 55 cm C18 column (1.9 µm particles, 100 Å pore size, ReproSil-Pur Basic C18, Dr. Maisch GmbH, Ammerbuch-Entringen, Germany) and eluted with a multi-step gradient, using buffer A (0.1% FA) and buffer B (80% ACN), from 2 to 7% buffer B in 5min, 7% to 23% buffer B in 70min, 23 to 45% buffer B in 30min and 45 to 95% buffer B in 5min, at a flow rate of 250 nL/min over 132 min. The column temperature was set to 60°C.

MS data were acquired using Xcalibur software using a data-dependent method. MS scans were acquired at a resolution of 70,000 and MS/MS scans (fixed first mass 100 m/z) at a resolution of 17,500. The AGC target and maximum injection time for the survey scans and the MS/MS scans were set to 3E6, 20ms, and 1E6, 60ms, respectively. An automatic selection of the 10 most intense precursor ions was activated (Top 10) with a 45s dynamic exclusion. The isolation window was set to 1.6 m/z and normalized collision energy fixed to 28 for HCD fragmentation. We used an underfill ratio of $1.00E^4$ for an intensity threshold of 1.7E5. Unassigned precursor ion charge states as well as 1, 7, 8, and >8 charged states were rejected, and peptide match was disable.

**Processing, quantification, and statistical analysis of MS data.** Raw data were analyzed using MaxQuant software version 1.5.5.1 (Cox and Mann, 2008) using the Andromeda search engine (Cox et al., 2011). The MS/MS spectra were searched against the mus musculus SwissProt database (53.449 entries from UniProt the 24/07/2018). Variable modifications (methionine oxidation, N-terminal acetylation) and fixed modification (cysteine carbamidomethylation) were set for the search and trypsin with a maximum of two missed cleavages was chosen for searching. The minimum peptide length was set to 7 amino acids and the false discovery rate (FDR) for peptide and protein identification was set to 0.01. At least a unique peptide per protein group was required for the identification of protein. The main search peptide tolerance was set to 4.5 ppm and to 20 ppm for the MS/MS match tolerance. Second peptides were enabled to identify co-fragmentation events and match between runs option for biological replicates of a same condition was selected with a match time window of 0.7 min for an alignment time window of 20 min. Quantification was performed using the XIC-based LFQ algorithm with the Fast LFQ mode as previously described (Cox et al., 2014). Unique and razor peptides, included modified peptides, with at least 2 ratio counts were accepted for quantification. For the differential analyses, proteins identified in the reverse and contaminant databases and proteins "only identified by site" were first discarded from the list of identified proteins. Then, only proteins with 6 quantified intensity values in a condition are kept. After log2 transformation, LFQ values were normalized by median centering within conditions (normalizeD function of the R package DAPAR) (Wieczorek et al., 2017). Remaining proteins without any LFQ value in one of both conditions have been considered as proteins quantitatively present in a condition and absent in another. They have therefore been set aside and considered as differentially abundant proteins. Next, missing values were imputed using the imp.norm function of the R package norm (Novo, 2013). Proteins with a fold-change under 2.0 have been considered not significantly differentially abundant. Statistical testing of the remaining proteins (having a fold-change over 2.0) was conducted using a limma t-test (Smyth, 2005) thanks to the R package *limma* (Ritchie et al., 2015). An adaptive Benjamini-Hochberg procedure was applied on the resulting p-values thanks to the function adjust.p of R package *cp4p* (Giai Gianetto et al., 2016) to estimate the proportion of true null hypotheses among the set of statistical tests. The proteins associated to an adjusted p-value inferior to an FDR of 1% have been considered as significantly differentially abundant proteins.

**Pathway and network analysis.** Gene ontology, pathway, and network analysis was performed using the GlueGO package, version 2.5.3, in Cytoscape (https://cytoscape.org/), version 3.7.1 [33,34]. Curated pathways for enrichment analysis were referenced from the database of GO Biological Function. For gene ontology data, testing was restricted to pathways with experimental evidence (EXP, IDA, IPI, IMP, IGI, IEP). The statistical cutoff for enriched pathways was Bonferroni-adjusted p-values <0.01 by right-sided hypergeometric testing. Pathway-connecting edges were drawn for kappa scores >40%. Kappa scores are a measure of inter-pathway agreement among observed proteins that indicate whether pathway agreement is greater than expected by chance based on shared proteins. Pathways with the same color indicate ≥50% similarity in terms. Pathway and network visualizations were generated and modified using the GlueGO package in Cytoscape.

**Immunohistochemistry and immunofluorescence.** TA muscles were isolated from mice and frozen directly in liquid nitrogen in cooled isopentane (320404, Sigma) for 40 seconds and Liver, pancreas, kidney and terotoma were imbedded in tissue freezing medium (14020108926, Leica), frozen in cooled isopentane on dry ice, all samples were stored at -80°C or directly cryosectioned in 10-$\mu$m sections.

For histology, sections were air dried and fixed with 4% PFA in PBS and washed, then stained with Haematoxylin and Eosin (H&E). For immunohistochemistry, the following primary antibodies were used: anti-Nanog (2ug/ml, H22, produced by Cambridge Research Biochemicals (Crb)). Slides were then incubated with the corresponding secondary antibodies conjugated with peroxidase from Dako (K4003).

**SAβGalactosidase (SAβGal) assay.** The assay was performed as previously described (Chiche et al., 2017). Briefly, sections were fixed at room temperature for 10 min in a solution containing 2% paraformaldehyde and 0.2% glutaraldehyde in PBS. Sections were washed in PBS and then incubated in an X-gal solution containing 4mM $K3Fe(CN)_6$, 4 mM $K4Fe(CN)_6$, 2 mM MgCl2, 0.02% NP-40 (Igepal) and 1mg/ml X-gal (EU0012, EUROMEDEX) in PBS pH = 5.5) at 37°C for 48 h. X-gal solution was changed after 24h. Samples were washed in PBS and post-fixed in 4% PFA in PBS for 15 min. After washes, samples were counterstained with Fast-red (H3403, VECTOR) and mounted in Eukitt (03989, Sigma). Finally, the sections were scanned using OLYPMPUS VS120, and SAβgal positive cells were quantified using self-developed program "Sendetect".

**BrdU immunofluorescence staining.** Cells were seeded on coverslip coated with Poly-L-Lysine (Sigma) in 24-well plates for analyses. Cells were incubated under different conditions for 72 hours then incubated for 1 hour with BrdU (10µM). Cells were washed with PBS and fixed in cold 70% EtOH (-20°C) for 20 minutes. Cells were washed with PBS and incubated with fresh 2N HCl for 20 min to denature DNA. Cells were washed 3 times with PBS and blocked into 3% BSA blocking solution for 30 min. After blocking cells were incubated with primary antibody for 2 hours at RT into humid chamber. Cells were washed with PBS three times prior incubation with secondary antibody for 45 min at RT. Finally, cells were mounted into DAPI Gold antifade mounting medium (Invitrogen, P36931). Images were acquired with an Olympus IX83 microscope and quantified using ImageJ software.

**Cell cycle analysis.** MEFs were incubated under different conditions. Briefly cells were incubated with BrdU (10µM) for 1h30 min then washed three times with PBS. Cells were collected and were washed with PBS, centrifuged and fixed in cold 70% EtOH. Cells were kept at -20°C until analysis. The day of analysis, fixed cells were centrifuged. Cells were washed with PBS and incubated with fresh 2N HCl for 20 min at RT. Cells were washed and incubated with FITC BrdU antibody (BD Pharmingen, 51-33284X) for 45 min at RT. Cells were washed with PBS, and resuspended in PBS containing Rnase A (100µg/mL) (Qiagen) and Propidium Iodide (50µg/mL) (Sigma Aldrich, P4864). Samples were analyzed at least 2 hours after using a Cytoflex Flow Cytometer (Beckman Coulter).

**Quantitative real-time PCR.** Total RNA was extracted from cells and tissue samples with Trizol (Invitrogen, 15596026) following provider's recommendations, samples were treated with DNase I before reverse transcription into cDNA following the manufacturer's protocol (Applied Biosystems, 4368813). Quantitative real-time PCR was performed using a LightCycler 480 (Roche) using SYBR Green Master (Roche). All values were obtained at least in duplicate, and in a total of at least two independent assays. Calculation for the values was made using the ΔΔCt method, as previously described (Yuan et al., 2006).

**Western Blot.** i4F MEFs were seeded on p60 plate and incubated under different conditions. Cells were collected at the indicated different timepoints directly in a mix composed of RIPA buffer (Sigma, R0278), DTT, Protease inhibitor cocktail (Sigma, 539131) and phosphatase inhibitor cocktail (Millipore, 524625). Samples were boiled for 5 min at 95°C before loading on polyacrylamide gels. Transfer occurred at 100V for 1hr on nitrocellulose membranes (BioRad). Membranes were blotted with TBST (0.2% Tween) and 5% BSA (Sigma, A3608) and incubated overnight with the primary antibody and for 45min with HRP-conjugated secondary antibody. Bands were revealed with Pierce ECL chemiluminescent substrate (Thermo Scientific, 32109) and analyzed using a ChemiDoc MP (BioRad).

**Primers**

| **Primers qPCR** | 5'- to -3' | |
|---|---|---|
| mGAPDH F | TTCACCACCATGGAGAAGGC | Li et al. 2009 |
| mGAPDH R | CCCTTTTGGCTCCACCCT | Li et al. 2009 |
| En-mOct4 F | TCTTTCCACCAGGCCCCCGGCTC | Abad et al., 2013 |
| En-mOct4 R | TGCGGGCGGACATGGGGAGATC C | Abad et al., 2013 |
| En-Sox2 F | TAGAGCTAGACTCCGGGCGATG A | Abad et al., 2013 |
| En-Sox2 R | TTGCCTTAAACAAGACCACGAA A | Abad et al., 2013 |
| En-Klf4 F | GCGAACTCACACAGGCGAGAAA CC | Abad et al., 2013 |
| En-Klf4 R | TCGCTTCCTCTTCCTCCGACACA | Abad et al., 2013 |
| Nanog F | AGGGTCTGCTACTGAGATGCTCT G | Abad et al., 2013 |
| Nanog R | CAACCACTGGTTTTTCTGCCACC G | Abad et al., 2013 |
| mLin28a F | GAAGAACATGCAGAAGCGAAGA | Li et al. 2009 |
| mLin28a R | CCGCAGTTGTAGCACCTGTCT | Li et al. 2009 |
| Occludin F | TGAAAGTCCACCTCCTTACAGA | https://pga.mgh.harvard.edu/primerbank/ |
| Occludin R | CCGGATAAAAAGAGTACGCTGG | https://pga.mgh.harvard.edu/primerbank/ |
| Zeb1 F | ACCGCCGTCATTTATCCTGAG | https://pga.mgh.harvard.edu/primerbank/ |
| Zeb1 R | CATCTGGTGTTCCGTTTTCATCA | https://pga.mgh.harvard.edu/primerbank/ |
| Zeb2 F | CCACGCAGTGAGCATCGAA | https://pga.mgh.harvard.edu/primerbank/ |
| Zeb2 R | CAGGTGGCAGGTCATTTTCTT | https://pga.mgh.harvard.edu/primerbank/ |
| Cdh1 (E-Cadherin)-F | CAGCCTTCTTTTCGGAAGACT | Li et al. 2009 |
| Cdh1 (E-Cadherin)-R | GGTAGACAGCTCCCTATGACTG | Li et al. 2009 |
| Amphiregulin F | GGTCTTAGGCTCAGGCCATTA | https://pga.mgh.harvard.edu/primerbank/ |

| | | |
|---|---|---|
| Amphiregulin R | CGCTTATGGTGGAAACCTCTC | https://pga.mgh.harvard.edu/primerbank/ |
| Betacellulin F | AATTCTCCACTGTGTGGTAGCA | https://pga.mgh.harvard.edu/primerbank/ |
| Betacellulin R | GGTTTTCACTTTCTGTCTAGGGG | https://pga.mgh.harvard.edu/primerbank/ |
| EGF F | AGCATCTCTCGGATTGACCCA | https://pga.mgh.harvard.edu/primerbank/ |
| EGF R | CCTGTCCCGTTAAGGAAAACTCT | https://pga.mgh.harvard.edu/primerbank/ |
| Epiregulin F | CTGCCTCTTGGGTCTTGACG | https://pga.mgh.harvard.edu/primerbank/ |
| Epiregulin R | GCGGTACAGTTATCCTCGGATTC | https://pga.mgh.harvard.edu/primerbank/ |
| EPGN F | GGGGGTTCTGATAGCAGTCTG | https://pga.mgh.harvard.edu/primerbank/ |
| EPGN R | TCGGTGTTGTTAAATGTCCAGTT | https://pga.mgh.harvard.edu/primerbank/ |
| HBEGF F | CGGGGAGTGCAGATACCTG | https://pga.mgh.harvard.edu/primerbank/ |
| HBEGF R | TTCTCCACTGGTAGAGTCAGC | https://pga.mgh.harvard.edu/primerbank/ |
| Neuregulin F | ATGGAGATTTATCCCCCAGACA | https://pga.mgh.harvard.edu/primerbank/ |
| Neuregulin R | GTTGAGGCACCCTCTGAGAC | https://pga.mgh.harvard.edu/primerbank/ |
| Neuregulin 2 F | TCGACCCTAACGGCAAAAACA | https://pga.mgh.harvard.edu/primerbank/ |
| Neuregulin 2 R | AACCAGCGATAGGAGGGCT | https://pga.mgh.harvard.edu/primerbank/ |
| Neuregulin 3 F | TTACGCTGTAGCGACTGCATC | https://pga.mgh.harvard.edu/primerbank/ |
| Neuregulin 3 R | GCCTACCACGATCCATTTAAGC | https://pga.mgh.harvard.edu/primerbank/ |
| Neuregulin 4 F | CACGCTGCGAAGAGGTTTTTC | https://pga.mgh.harvard.edu/primerbank/ |
| Neuregulin 4 R | CGCGATGGTAAGAGTGAGGA | https://pga.mgh.harvard.edu/primerbank/ |
| TGFa F | CACTCTGGGTACGTGGGTG | https://pga.mgh.harvard.edu/primerbank/ |
| TGFa R | CACAGGTGATAATGAGGACAGC | https://pga.mgh.harvard.edu/primerbank/ |
| TGFb F | CACTCTGGGTACGTGGGTG | https://pga.mgh.harvard.edu/primerbank/ |
| TGFb R | CACAGGTGATAATGAGGACAGC | https://pga.mgh.harvard.edu/primerbank/ |
| p16 INK4A F | CGTACCCCGATTCAGGTGAT | Li et al. 2009 |
| p16 INK4A R | TTGAGCAGAAGAGCTGCTACGT | Li et al. 2009 |
| ARF F | GCCGCACCGGAATCCT | Li et al. 2009 |
| ARF R | TTGAGCAGAAGAGCTGCTACGT | Li et al. 2009 |
| mIL-6 F | CTGCAAGAGACTTCCATCCAG | https://pga.mgh.harvard.edu/primerbank/ |

| | | |
|---|---|---|
| mIL-6 R | AGTGGTATAGACAGGTCTGTTGG | https://pga.mgh.harvard.edu/primerbank/ |
| MMP3 F | CCTGATGTTGGTGGCTTCA | https://pga.mgh.harvard.edu/primerbank/ |
| MMP3 R | TCCTGTAGGTGATGTGGGATTTC | https://pga.mgh.harvard.edu/primerbank/ |
| MMP13 F | ACTTCTACCCATTTGATGGACCTT | https://pga.mgh.harvard.edu/primerbank/ |
| MMP13 R | AAGCTCATGGGCAGCAACA | https://pga.mgh.harvard.edu/primerbank/ |
